# Development and Clinical Evaluation of a Multiplexed Health Surveillance Panel Using Ultra High-Throughput PRM-MS in an Inflammatory Bowel Disease Cohort

**DOI:** 10.1101/2025.04.02.646850

**Authors:** Qin Fu, Philip M. Remes, Jihyeon Lee, Cristina Jacob, Dalin Li, Manasa Vegesna, Koen Raedschelders, Ali Haghani, Emebet Mengesha, Philip Debbas, Estelle Hoedt, Sandy Joung, Susan Cheng, Scott Peterman, Justyna Fert-Bober, Gil Y. Melmed, Dermot P.B. McGovern, Christopher I. Murray, Jennifer E. Van Eyk

## Abstract

Despite advances in clinical proteomics, translating protein biomarker discoveries into clinical use remains challenging due to the technical complexity of the validation process. Targeted MS-based proteomics approaches such as parallel reaction monitoring (PRM) offer sensitive and specific assays for biomarker translation. In this study, we developed a multiplex PRM assay using the Stellar mass spectrometry platform to quantify 57 plasma proteins, including 21 FDA-approved proteins. Loading curves (11-points) were performed at 4 sample throughputs (100, 144, 180, and 300 samples per day) using independent, optimized, and scheduled PRM methods. Following optimization, an inflammatory bowel disease (IBD) cohort of plasma samples (493 IBD, 509 matched controls) was analyzed at a throughput of 180 SPD. To monitor system performance, the study also included 1,000 additional injections for system suitability tests, low-, middle-, and high-quality controls, washes, and blanks. Using this approach, we observed high quantifiability (linearity, sensitivity, reproducibility) in the PRM assay and consistent in data acquisition across a large cohort. We also validated the candidate IBD markers, C-reactive protein and orosomucoid protein, identified in a recent discovery experiment.

## Introduction

Blood plasma is a readily accessible biofluid that reflects the physiological state of an individual, making it an invaluable source for biomarkers of disease (1). After a potential biomarker has been identified in a discovery experiment, thorough validation in large patient cohorts is necessary to assess a biomarker’s efficacy and clinical utility. One option for validation is a mass spectrometry (MS)-based targeted protein assay where a peptide (precursor) representing the candidate protein is quantified by measuring its intensity across the chromatographic peak during elution. Each targeted assay must be precise, reliable, robust over repeated measurements, and scalable to the high sample throughputs necessary for large scale analysis (2-5). An advantage of a targeted MS protein assay is that individual protein assays can be combined, or multiplexed, allowing several candidate biomarkers can be quantified simultaneously. Combining the analysis significantly reduces instrument time during the lengthy validation phase, however the development of high-quality targeted multiplex assays remains challenging. Each analyte must be optimized and scheduled with respect to the others resulting in a time-consuming process that can delay translation. Currently, there is no single solution to address the scale, sensitivity, and throughput demands of a multiplexed assay that can meet the rigor of clinical validation.

Targeted MS-based proteomics approaches, such as multiple reaction monitoring (MRM) and parallel reaction monitoring (PRM), have been deployed as single or multiplexed assays to validate candidate biomarkers (6, 7). For each approach, a representative peptide precursor is targeted by the instrument and quantified. MRM assays are traditionally performed using a triple quadrupole mass spectrometer. In an MRM assay, the desired precursor is isolated in the first quadrupole (Q1), fragmented in the second (Q2) via collision-induced dissociation and verified by monitoring multiple predefined fragments ions, also referred to as transitions in the third quadrupole (Q3). The MRM triple quadrupole approach excels at targeted detection however it requires defining MRM transitions pre-experiment, since triples are not well-suited for acquisitions that scan over a wide m/z range (8). PRM assays differ in that all fragment ions from a targeted precursor are accumulated in parallel and measured in a single acquisition event. The parallel accumulation gives PRM a sensitivity advantage over MRM; it also greatly simplifies the experimental definition and effectively increases selectivity because transitions can be defined post-experiment. PRM assays have traditionally been performed on Quadrupole-Orbitrap or Quadrupole-Time of Flight (TOF) mass spectrometry systems. Quadrupole-TOF systems are fast, and the latest generation is sensitive, while generally offering lower resolution and mass accuracy compared to Quadrupole-Orbitrap mass spectrometers. Quadrupole-Orbitrap mass spectrometers conversely, provide higher resolution and mass accuracy, but have lower acquisition rates and use a less sensitive image-charge detection mode compared to the faster triple quadrupole and TOF instruments which use electron-multiplier-based detectors (9, 10). A recent innovation has introduced a dual-pressure linear ion trap in the Q3 position of the triple quadrupole design (11). Replacing the third quadrupole with a linear ion trap creates a hybrid instrument with triple quadrupole-like speed and the PRM sensitivity advantage, albeit with unit mass resolution and accuracy. This new instrument configuration allows for improved acquisition rate (up to 140 Hz), greater ion storage, higher mass resolution, faster cycle time, extended dynamic range and MS^n^ acquisition (11). This MS platform also uses the Thermo Scientific™ Adaptive RT algorithm to adjust acquisition windows in real-time to account for drift, and the Skyline-based tool, PRM Conductor, that automates the selection and dynamic scheduling of precursors, simplifying the assay development process (11).

To test the hybrid configuration, we re-designed our previously reported Health Surveillance Protein (HSP) MRM assay as a multiplexed PRM assay (12, 13). The HSP was created as a diagnostic tool to measure 57 laboratory developed tests (LDT) biomarkers related to cardiovascular, inflammatory, and metabolic conditions. The assay leverages 21 already FDA-approved biomarkers which are routinely measure clinically and included C-reactive protein (CRP). The HSP assay uses 83 stable isotopically labeled (SIL) peptides representing the 57 proteins to provide precise quantitation. First, we tested the HSP PRM assay using an 11-point dilution series at 4 different sample throughputs to find the optimal balance in speed and quantification of all proteins. Then we applied this re-tooled assay to a cohort of inflammatory bowel disease (IBD) patients. IBD, comprising Crohn’s disease (CD) and ulcerative colitis (UC), is a complex immune-mediated disorder influenced by genetic predispositions and environmental factors (14). While the mechanisms underlying IBD pathogenesis and progression have been extensively studied, significant gaps remain in understanding the drivers of disease variability and therapeutic responses. We have previously analyzed a cohort of IBD patients and age- and sex-matched healthy subjects in a biomarker discovery study where we identified C-reactive protein (CRP) and orosomucoid protein (A1AG1) as candidates (15). CRP is currently used clinically to assess disease activity in Crohn’s disease and to a lesser degree for ulcerative colitis, where levels of circulating CRP indicate overall inflammatory activity, severity of flare, or response to therapy (16-18). A1AG1 is also an acute phase protein and reflects similar responses to CRP (19).

Using this new MS-platform we validate the candidate IBD biomarkers and assess the performance of a large-scale multiplex PRM assay in over 1000 clinical case and control patients. We also compared CRP measured in the clinical chemistry laboratory as part of standard care for the IBD patients with the CRP measured within the HSP PRM assay and found a correlation of 0.907 between the two independent assays. This work will establish a robust framework for precision and scalability in biomarker validation and could be broadly applicable to other translational studies and disease contexts requiring precise relative quantification of extensive protein sets.

## Results and Discussion

### PRM Method Development

We evaluated a modified 57-protein multiplexed HSP PRM method on a high-speed hybrid nominal mass platform (Stellar-MS, Thermo Fisher Scientific). This modified 57-protein multiplexed HSP PRM method was developed based on a previously published MRM assay (12, 20). The current iteration of the assay consists of 83 prototypic peptides, their corresponding SIL peptides, and a 14 Peptide Retention Time Calibration (PRTC) mixture (Thermo Fisher Scientific, Rockford, IL), totaling 180 peptides. To develop a large scale and high-throughput PRM assay, we assessed four different throughputs: 100, 144, 180, and 300 samples per day (SPD). For large-scale studies, minimizing the acquisition time is essential for efficient use of the instrumentation. Method development is a multi-step process, as illustrated in Figure 1. Methods for each sample throughput were developed using Skyline with Prosit to predict the spectral library, precursors and indexed retention times (iRTs) necessary to generate an unscheduled PRM method for the 83 different peptide sequences (21-24). Next, the instrument parameters for each unscheduled method were optimized using PRM Conductor (PMID: 38854069) and Skyline using the results from replicate injections of the 83 SIL peptides in solvent (0.1% FA) and SIL peptides spiked into 300 ng of digested plasma and with the PRTC mixture (11). The throughput-dependent PRM conductor parameters are summarized in Supplemental Table 1; the LC conditions, peptide retention times, precursor m/z, charges are detailed in Supplemental Table 2 and 3.

**Figure 1.**
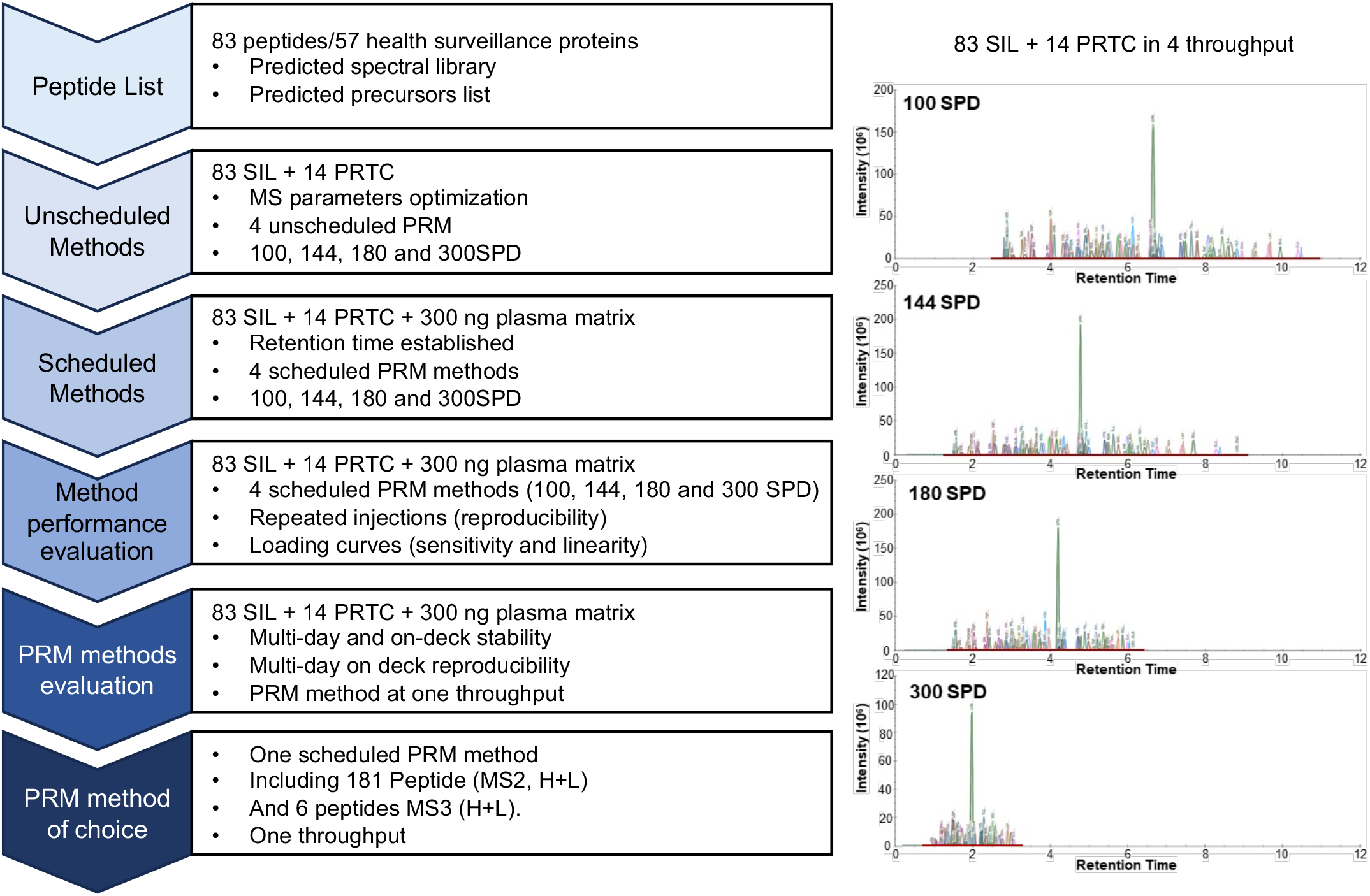
Schema of scheduled PRM methods development, evaluation, and establishment of method of choice for large scale and high-throughput analysis.

### Evaluation of Four Throughput Methods for Reproducibility, Sensitivity, and Linearity

The performance of the four different throughput methods (100, 144, 180, and 300 SPD) was evaluated using an 11-point SIL peptide dilution curve spiked into a pooled human plasma sample from 100 healthy females and 100 healthy males. As a benchmark, the time for 10 injections is shown for each throughput method (Figure 2A). For each dilution curve, points per peak, peak area, and coefficient of variation (CV) were assessed for all 83 peptides (Figure 2 B-D). As expected, the median number of points per precursor peak decreased with increasing sample throughput. Only the 300 SPD method failed to meet the required 7 points per peak for accurate quantification and precision (25). Furthermore, the median peak areas also decreased with increasing sample throughput. The median CV values were all <=10%, indicating good reproducibility across all four methods.

**Figure 2.**
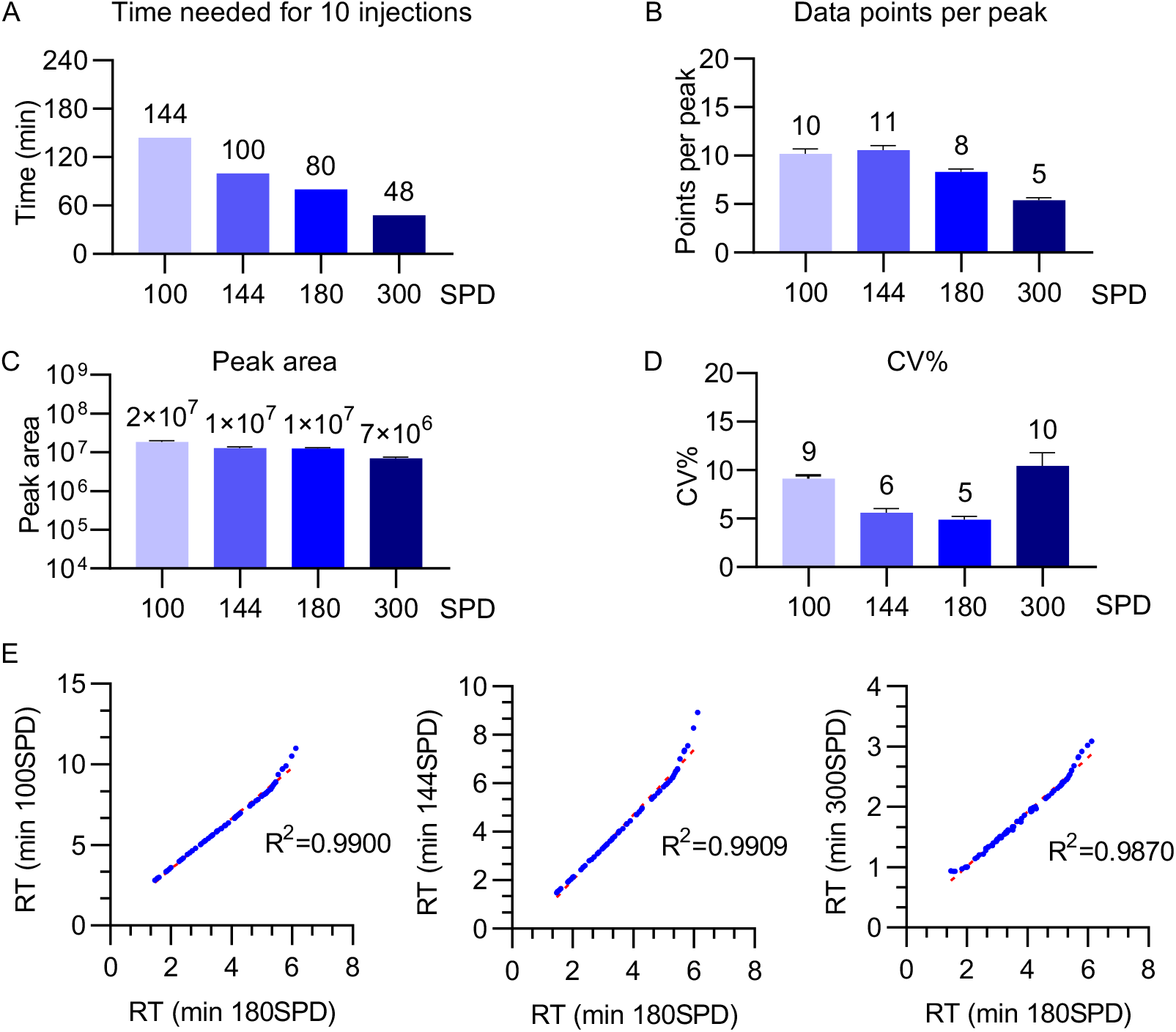
Evaluation of four PRM methods with repeated injections. The performance of the 4 SPD methods was initially evaluated using 30 fmol of the 83 SIL peptides spiked into 300 ng of digested control plasma (n=10, technical replicates). **A**. Indicates the time required for 10 injections with each method. **B**. The median cross-peak points. **C**. The median peak intensities. **D**. The median coefficient of variation (CV) of the signal intensities. **E**. Correlation between the retention times of 83 SIL across any two throughput conditions.

A key aspect of developing a multiplexed PRM assay is consistency in retention time to conduct effective scheduling of multiple precursors. We examined the retention time correlation across all four throughput settings for 83 SIL. Pairwise correlation analysis between any two throughput settings demonstrated excellent correlation, with R^2^ values exceeding 0.98 (Figure 2E).

Next, the sensitivity, linearity, LLOD (lower limit of detection), LLOQ (lower limit of quantification), and reproducibility for each of the 83 peptides was determined across the 11-point dilution curves for each SPD method (Figure 3). The observed median sensitivity for the 83 SIL peptides across all the methods was sub-fmol (LOD < 0.5 fmol), with the longer SPD methods achieving a higher percentage of peptides at or below this level (Figure 3A). The performance of the LLOQ (<5 fmol) was similar among 100, 144 and 180 SPD. There was a significant drop-off at the 300 SPD throughput (81-82% for the lower SPDs vs 62% for 300 SPD) (Figure 3B). The linear response of each peptide was also assessed, indicating that more than 95% of all peptides demonstrated an R^2^>0.9 across the 11-point dilution curve (Figure 3C) for all the SPD methods. Using the A2GL; ALGHLDLSGNR SIL peptide as an example, Figure 3D shows SIL standard curves across four different SPDs, along with LLOD, LLOQ, R^2^ values, and precursor chromatographs. The asymmetric and imperfect chromatograms observed for this peptide at 300 SPD are consistent with low data points per peak and a relatively high LOQ. The LLOQ, LLOD, and R^2^ for each peptide at all four SPDs are summarized in Supplement Table 4. Based on the initial characterization of the methods, the 180 SPD method was found to have the preferred balance of performance and sample throughput. All subsequent evaluations were performed using this method.

**Figure 3.**
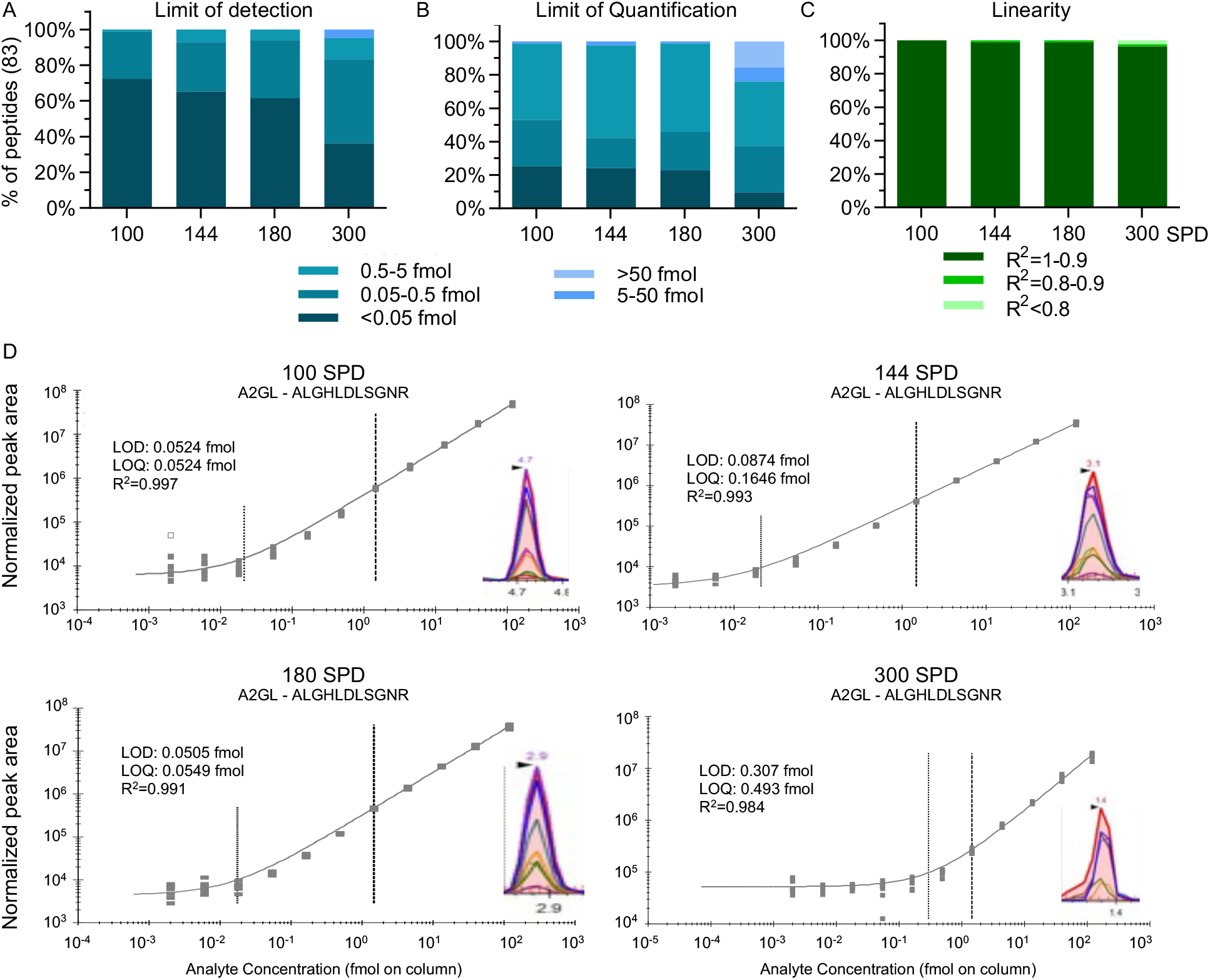
Evaluation of the sensitivity and linearity of four PRM throughput methods. 0.002-120 fmol 83 SIL/300 ng pooled plasma peptides was evaluated for each SPD method (11-point dilution curve; n=5, technical replicates). **A**. The LLOD was calculated using the Skyline bilinear regression fit (blank+3x standard deviation). The results are binned into 5 ranges. **B**. The LLOQ was also determined as coefficient of variation (CV) <20% and a bias <20%. The results are binned into the same 5 ranges. **C**. Median linear response (R^2^) for all 83 SIL peptides over the dilution curve. **D**. Examples of linear regression curves for A2GL_HUMAN and ALGHLDLSGNR are presented for 100, 144, 180, and 300 SPD. The peaks displayed are from Skyline showing integrated peaks of the same peptides.

### PRM Assay Reproducibility

The precision of the PRM assay on the Stellar MS platform was assessed for within-sample and across-day repeatability using a 5×5×5 experiment (26). Five injections per day over 5 days were performed using a number of independently pooled human plasma samples (labeled as Pools A-E, with each pool consisting of eight different plasma samples) spiked with the 83 SIL peptide panel (Figure 4A). The mean intra-assay (same day, between pools), inter-assay (same pool, between days) and total (all pools, all days) coefficients of variation (CV) were calculated from the area ratio of the endogenous and SIL peptides (Light/Heavy ratio; L/H) (Figure 4B-D) (See Supplemental Table 5). For the intra-day precision of the 83 peptides in the 5 different plasma pools tested, 74-76 of the peptides had intra-day mean CV of <10%, with only 3-4 peptides >20%. The evaluation of between-day precision revealed that 78-80 peptides had inter-day mean CV of <10% with 1-4 >20%. Finally, across all 125 injections spanning the 5 days, 72-74 peptides had total experiment CVs of <10% and only 3-5 peptides had a CV >20%. The peptides observed with greater than 20% CV inter-, intra or total were due to low concentration of the native peptide in the plasma.

**Figure 4.**
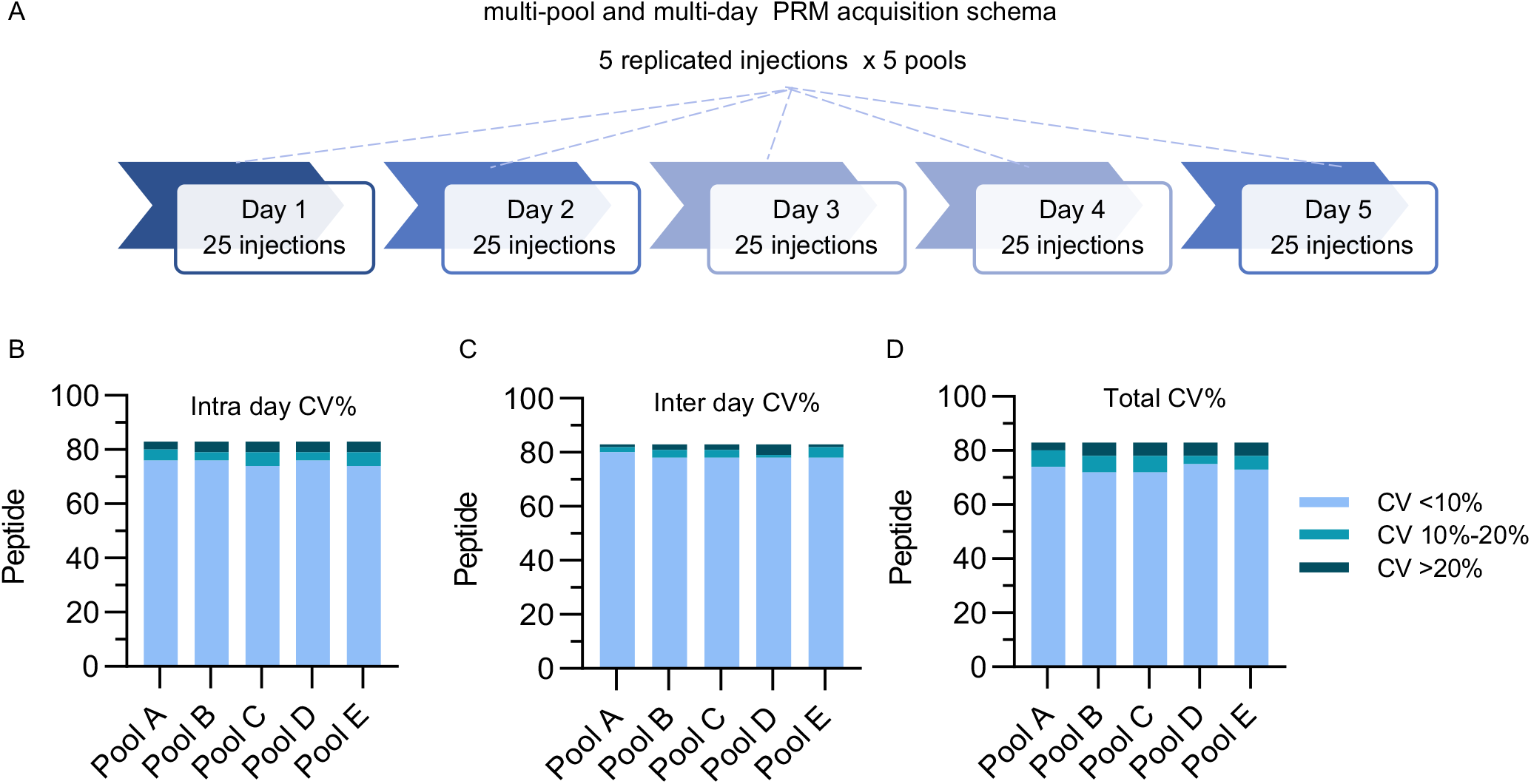
Evaluation of reproducibility. **A**. Schematic outlining the analysis of 5 unique pooled plasma samples collected over 5 different days using the 180 SPD method (n=5, biological; n=5, technical). Each plasma pool was created by combining eight individual plasma samples; a total of 40 samples used to generate five distinct plasma pools. Summary of the median intra-day (**B**) and inter-day (**C**) CV for all 83 SIL peptides. **D**. Total variability was assessed using the mean intra-day CV (across five days) and the mean inter-day CV (across all five replicates) with the following formula: CV total = (CV intra^2^ +CV inter^2^)^1/2^. The total median variability is reported for each pool.

### PRM Data Acquisition of the IBD Patient Cohort

As a proof of principle for the platform, we evaluated the application of the HSP 57-protein multiplex PRM assay in a clinical cohort. We assembled an IBD cohort that comprised 1002 individual plasma samples (n=493 IBD patients and n=509 age- and sex-matched healthy control subjects). The full cohort was processed across twelve 96-well plates using an i7 automated workstation (Beckman Coulter). The i7 automation workstation was previously set up for routine bottom-up sample preparation, which includes protein denaturation, Cys reduction and alkylation, trypsin digestion and desalting (27). After desalting, the SIL peptides were spiked in prior to LC-MS PRM analysis. LC-MS acquisition was completed over 12 days at 180 SPD. The PRM acquisition sequence schema was designed to maximize instrument utilization by continuously acquiring data and simultaneously evaluating LC-MS system performance. Throughout a 96-well sample plate, plasma sample injections were intermixed with an equal number of injections for quality assurance (Figure 5). Column washes and blank solvent injections were performed every 8 samples to assess column carry-over. Patient samples were run in blocks of 4 with a PRTC blank acquired before and after each block. Before and after every 2 patient blocks, a system suitability test (SST) was performed. The SST samples comprising 300 ng pooled plasma peptides, 14 PRTC peptides and 150 fmol 83 SIL peptides were used to assess signal stability and/or drift in retention time. Finally, at the beginning, mid-point, and end of each plate, three quality control (QC) samples were run, with 300 ng of control plasma spiked with 83 SIL peptides at high (120 fmol), moderate (40 fmol) and low (5 fmol) concentrations to monitor instrument performance.

**Figure 5.**
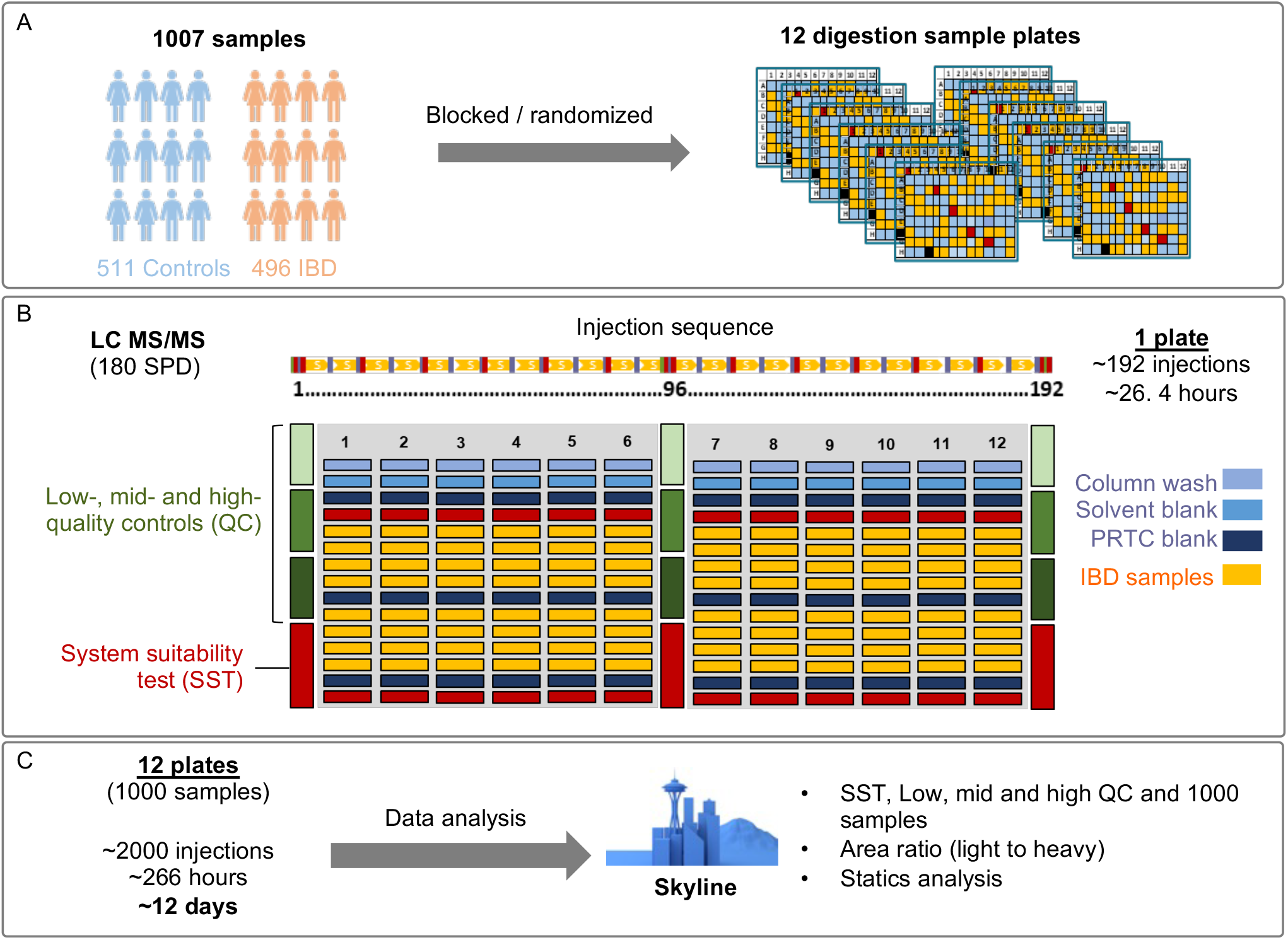
Design of 57-Plex PRM-MS clinical cohort analysis. **A**. Schema for processing and assignment of randomized blocks of subject plasma samples in 12 96-well plates for analysis. **B**. Schematic of the plate layout for the patient and QC samples. Each plate included LC column washes, solvent banks, and PRTC blanks for every IBD cohort 8 samples. System suitability tests were monitored for every four samples. Quality controls were implemented at three levels: High, mid and low (120, 40 or 5 fmol SIL/300 ng plasma), assessed at the beginning, middle, and end of each plate. The complete acquisition process for 1,002 samples (12 plates) took over 12 days and >2000 injections. **C**. PRM peak integration and data analysis for the SST, QC, and patient samples were performed using Skyline.

In total, 311 SST runs were completed during the acquisition of the entire 1002-sample IBD cohort dataset. The retention times of the 83 spiked SIL peptides demonstrated good stability. On average, the first peptide (VVLSQGSK, SHBG) eluted at 1.474±0.015 minutes, whereas the last peptide (SLEYLDLSFNQIAR, LUM) eluted at 6.177±0.039 minutes (Figure 6A). The median CV for the retention times was 3%, indicating strong stability across all 83 peptides. The total peak area of the 83 heavy peptides had a CV of 26%, further indicating the reproducibility of the SIL signal throughout the data acquisition process (Figure 6B). The evaluation of the QC samples showed good consistency across the cohorts. The three low-, middle-, and high-QC samples were collected at the beginning, middle, and end of each IBD sample plate, resulting in a total of 108 QC samples across the 12-plate acquisition. We consistently observed reproducible signals for the 83 spiked SIL peptides in all the QC runs. The median CVs for these peptides were 23%, 22%, and 23% for high, middle, and low QCs, respectively (Figure 6C). The RT consistency for each of the 83 spiked SIL internal standard peptides was also investigated. The RT CV% ranged from <1% to 6% and the 83 unnormalized SIL intensity CV% ranged from 13% to 55% across 1007 plasma samples (Figure 6D).

**Figure 6.**
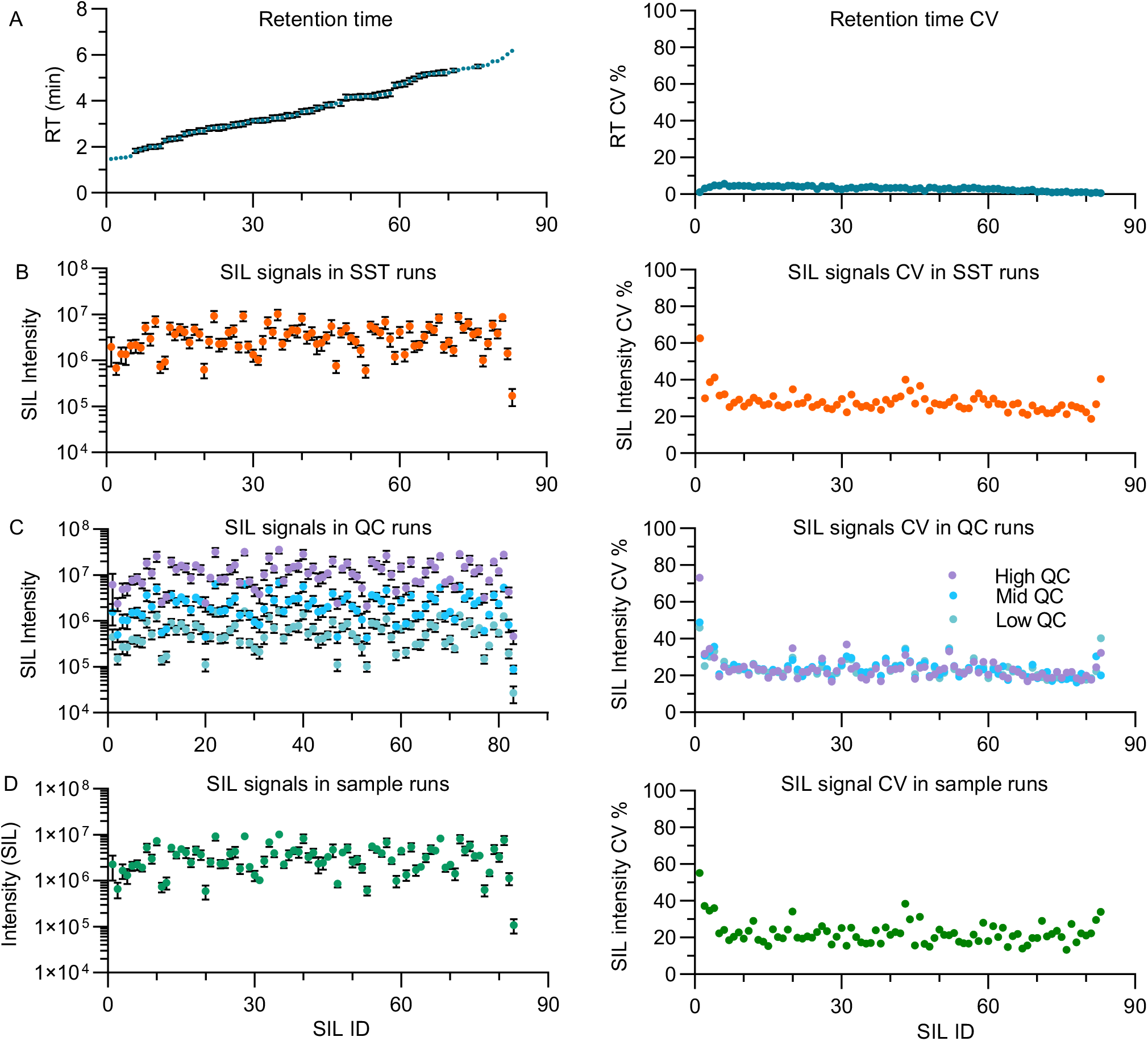
Quality assessment of PRM acquisition. **A**. Plot of the median RT (left) and CV (right) in the SST samples for the 83 SIL peptides (error bars = SD; n=331, technical). The SIL IDs were sorted based on retention time, arranged from low to high. **B**. Plot of the median SIL intensity (left) and CV (right) for the SST samples. **C**. Plot of the median SIL intensity (left) and CV (right) for the high (120 fmol), mid (40 fmol) and low (5 fmol) QC samples (error bars = SD; n=36 each load, technical). **D**. Plot of the median SIL intensity (left) and CV (right) for the patient samples (error bars = SD; n=1002, biological).

### LC-MS^3^ Strategy Increases the Sensitivity and Specificity

One option to potentially increase selectivity and sensitivity is the use of MS^3^ to the further isolate, fragment, and detect product ions.(28) We compared the MS^2^ and MS^3^ strategies for the CRP peptide (ESDTSYVSLK) in the IBD cohort (Figure 7A). We observed that the targeted MS^3^ method demonstrated a 9-fold increase in sensitivity, with a lower limit of quantification (LLOQ) of 0.0183 fmol/mL for MS^3^ compared with 0.1646 fmol/mL for MS^2^. The CRP MS^3^ method was subsequently incorporated into the final PRM assay method for the panel of 57 HSP panel assay. The 9-fold improvement in sensitivity for detecting CRP highlights the platform’s ability to identify low-abundance peptides effectively. Furthermore, by targeting specific ion fragments, MS^3^ minimizes interference from co-eluting or isobaric peptides, which is crucial for complex matrices like plasma or serum.

**Figure 7.**
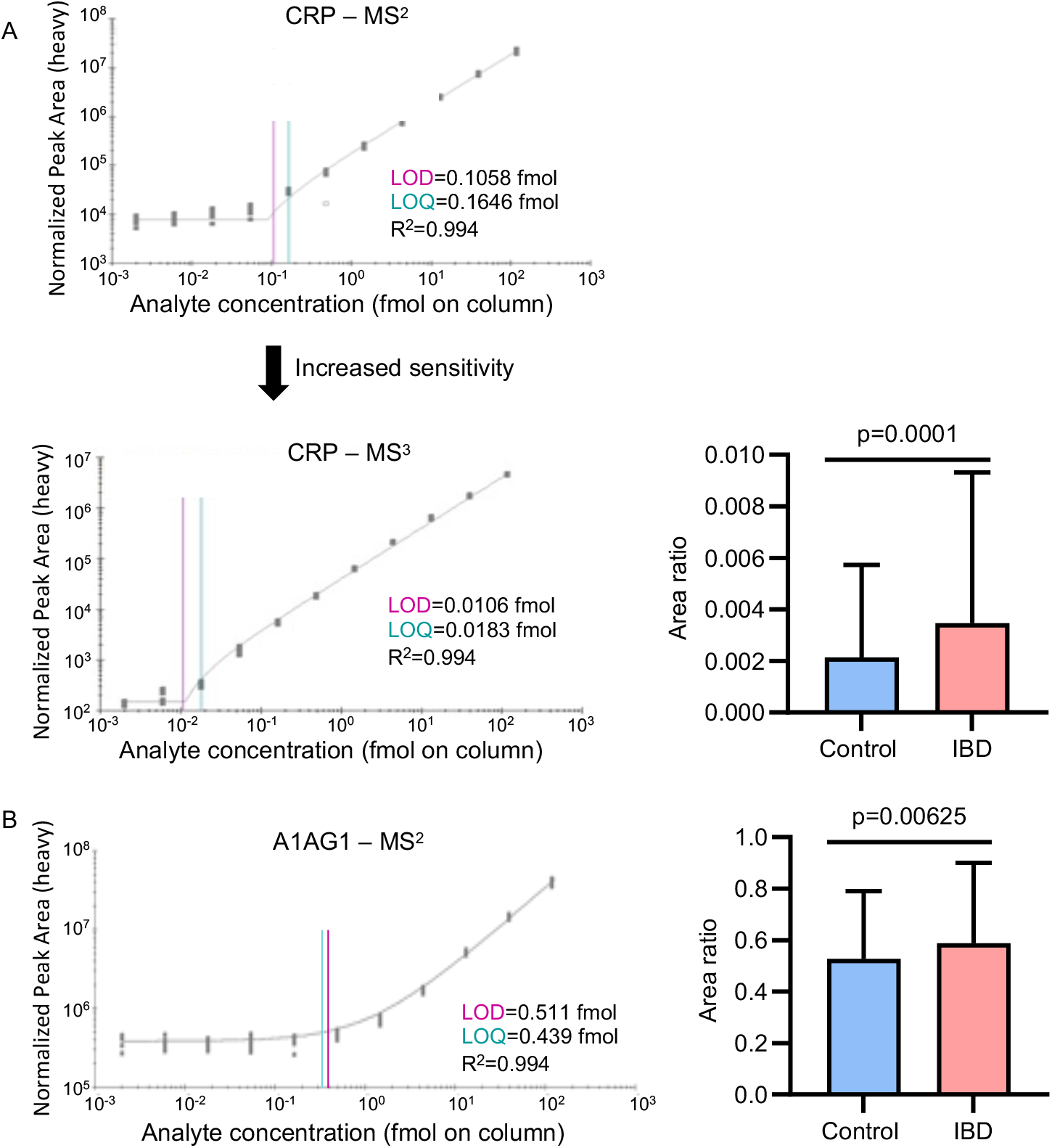
Improved sensitivity of detection and validation of candidate IBD biomarkers. **A**. PRM measurement of CRP precursor ESDTSYVSLK over the dilution series (11-points) using MS^2^ (upper) and MS^3^ (lower) detection to measure LOD (purple line) and LOQ (green line) for each approach (n=5, technical). The MS^3^ approach was used to analyze the IBD cohort. CRP was significantly increased in IBD patients compared to control patients (lower right). **B**. PRM detection of A1AG1 precursor SDVVYTDWK in MS^2^ mode over the dilution series (11-point) to measure LOD and LOQ (n=5, technical) (left). In cohort analysis, A1AG1 was significantly increased in IBD patients (right) (error bars = SD, n=496 IBD; n=511 control).

### Validation of IBD Biomarkers using the HSP PRM Assay

In our previously published IBD discovery research, CRP and A1AG1 were identified as significantly upregulated proteins in the IBD population (15). CRP is routinely measured as part of the diagnostic and monitoring procedures for IBD patients (29). A1AG1, also known as orosomucoid protein, has been associated with IBD (19). In our targeted PRM study, both CRP (ESDTSYVSLK) and A1AG1 (SDVVYTDWK) are included in our 57 protein-HSP panel. Consistent with the discovery results, our PRM analysis with internal standard analysis validated the CRP and A1AG1 results, both were found to be upregulated in the IBD population, (P=0.0001 for CRP and P=0.00625 for A1AG1) (Figure 7).

We further confirmed the CRP measured with PRM-MS by comparing it to CRP measured from the IBD patients using an antibody-based clinical assay. CRP is frequently measured as part of the diagnostic and monitoring procedures for IBD patients and was independently measured in 91 IBD patients at the same time point of sample collection (42 at 8 weeks, 44 at 16 weeks and 5 at 24 weeks) in a clinical lab using standard procedures (30). We examined the correlation between the CRP level measured by PRM and the clinically measured CRP level. We observed a strong correlation between CRP levels measured by MS and a standard clinical test, with a spearman correlation of 0.907 (Figure 8). Our results demonstrated validity of the MS-based and high throughput PRM analysis.

**Figure 8.**
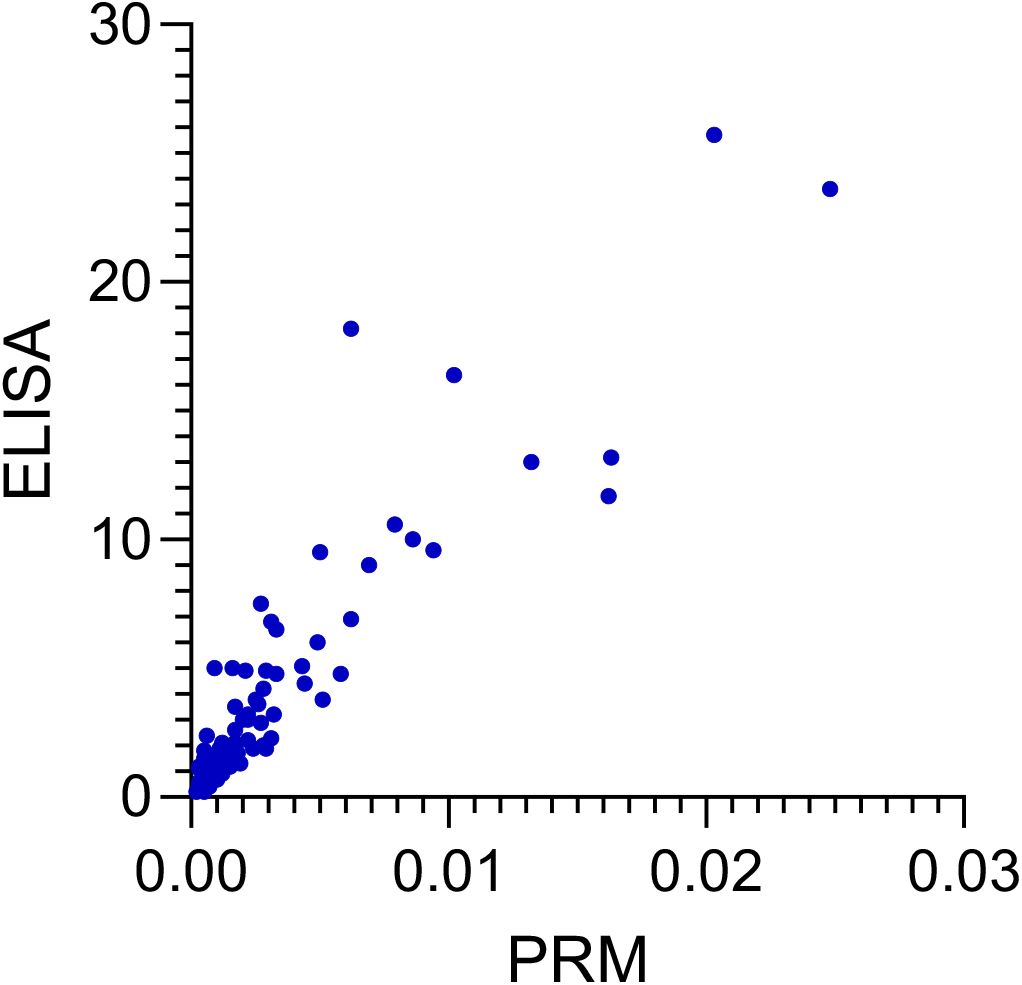
Comparison of PRM with ELISA measurements for CRP. Plot of the correlation of CRP PRM-MS based analysis (area ratio of light/heavy) and antibody-based ELISA assay (ng/mL) in IBD patient plasma (n=91). A Spearman correlation test revealed a strong correlation between the antibody-based ELISA assay and the mass spectrometry based PRM assay (r = 0.907, p (two tailed) <0.0001, 95% confidence interval 0.8603 to 0.9384), which was statistically significant.

## Conclusion

The final multiplex PRM assay offers enhanced sensitivity without compromising throughput, making it a powerful tool for the accurate quantification of multiple biomarkers and translational research, including IBD research, enabling the identification and quantification of a panel of biomarkers with high precision. Future work should focus on further refining the assay for broader clinical applications, including other inflammatory and chronic diseases. Additionally, the scalability of this platform offers opportunities for longitudinal studies and real-time monitoring of disease progression, paving the way for precision medicine approaches in clinical proteomics. This work will establish a robust framework for precision and scalability in biomarker validation and could be broadly applicable to other translational studies and disease contexts requiring precise relative quantification of extensive protein sets. Overall, these results validate the feasibility of the multiplex PRM assay for high-throughput proteomic analyses, with robust reproducibility, stability, and performance across a large clinical cohort.

## Supporting information

STable1

STable2

STable3

STable4

STable5

STable6

## Ethics

Ethical approval to obtain human samples and perform the research outlined was obtained from the Institutional Review Board of Cedars-Sinai Medical Center on Research Involving Human Subjects (Cedars-Sinai Medical Center IRB#: Study 00001411 and Study 00000621). The study was conducted in accordance with the ethical principles specified in the Declaration of Helsinki. Informed consent was obtained from all participants.

## Data Availability

The mass spectrometry proteomics raw data files have been deposited to the The ProteomeXchange ID reserved for the data is: PXD062364. Data will be made available to the public upon publication.

## Acknowledgements

The authors wish to express their gratitude for the contributions that have greatly facilitated the success of this study. Special thanks are extended to Saeed Seyedmohammad and Lindsey Becker for their assistance in sample organization and aliquoting as well as Maxim Zhgamadze, Nathan Hendricks, and Angel Keoseyan for sample preparation.

This study was supported by 1 U01 DK124019-01 (JVE), 1 R01 HL155346-01A1 (JVE), the Leona M. and Harry B. Helmsley Charitable Trust, the Fred L. Hartley Family Foundation, the Widjaja Foundation Inflammatory Bowel and Immunobiology Research Institute, National Institute of Diabetes and Digestive and Kidney Disease Grant U01DK062413 (DPBM), the Cedars-Sinai Precision Health Initiative, and the Erika J. Glazer Family Foundation.

