## Supplemental Methods for "Development and Clinical Evaluation of a Multiplexed Health Surveillance Panel Using Ultra High-Throughput PRM-MS in an Inflammatory Bowel Disease Cohort"

### Experimental Methods

#### Plasma Samples.

Pooled human plasma from 100 healthy females and 100 healthy males ($K_2EDTA$, 0.2 µm filtered) was obtained from BioIVT (Westbury, NY) and used for all method development, assay performance evaluation, loading curves, controls, and system suitability tests. This same pooled plasma sample has been used previously (1, 2).

#### IBD and Control Cohorts.

Details of the IBD and control cohorts have been described previously (3, 4). In brief, study subjects were enrolled in an IRB-approved prospective registry at Cedars-Sinai Medical Center (CSMC), and plasma samples were collected between December 2020 and June 2021. The Corale IBD cohort consisted of 493 subjects aged 13 years and older with a confirmed IBD diagnosis based on standard clinical criteria, and the Healthcare Worker (HCW) cohort consisted of 509 healthcare professionals at the CSMC. Both cohorts were originally designed to study the effects of SARS-CoV-2 vaccines in parallel and participants in both cohorts received two doses of the mRNA vaccine per the local department of health guidelines and institutional policies. The samples were collected at 8, 16, and 24 weeks after the 2nd dose of the COVID-19 vaccine in both cohorts. Supplemental table 6 lists the sex, age, and clinical designation of the IBD patients and control subjects. All human plasma was aliquoted and stored at -80$^{o}$C until processing.

#### Automated Plasma Sample Digestion and Desalting.

Automated digestion and desalting of the plasma samples were performed as previously described (5). The plasma samples (5 µL) were processed via the BioMek i7 automated workstation. The samples were denatured in a solution of 350 mL/L 2,2,2-trifluoroethanol (TFE) and 40 mM dithiothreitol in 50 mM $NH_4CO_3$ at 60°C for one hour and alkylated with 10 mM iodoacetamide at room temperature (25°C) for 30 minutes. The reactions were quenched by adding 5 mM dithiothreitol at 25°C for 15 minutes. The samples were then diluted with 50 mM $NH_4CO_3$, digested with trypsin at a protein-to-enzyme ratio of 25:1, and incubated for 4 hours at 37°C. The digestion was stopped by adding formic acid to achieve a final concentration of 2%. Desalting was performed via a positive-pressure apparatus and an i7 Biomek workstation. For desalting, 300 µL of the digested sample was mixed with 850 µL of a solution containing 2% phosphoric acid and 0.1% formic acid and applied to an Oasis 30 µm HLB 96-well plate (Waters Co.). Following activation with 1 mL of methanol and washing with 3 × 1 mL of 0.1% formic acid, 1150 µL of each sample was loaded onto an HLB plate and eluted with 1 mL of a 50% acetonitrile solution containing 0.1% formic acid. The samples were evaporated to dryness and stored at -80°C until further analysis. At the time of MS analysis, the peptides were resuspended in a 0.1% formic acid solution, and iRT standards were added prior to the LC–MS/MS run.

**LC–MS/MS Analysis.**

A Vanquish Neo UHPLC system (Thermo Fisher Scientific) was coupled with a Stellar mass spectrometer (Thermo Fisher Scientific) featuring an EASY-spray source (Thermo Fisher Scientific) for data acquisition. Ionization was achieved via an Easy-Spray™ source at 2000 V, with the ion transfer tube maintained at 275°C. The digested plasma samples were separated over total run times of 14 minutes (100 SPD), 10 minutes (144 SPD), 8 minutes (180 SPD), and 4.8 minutes (300 SPD) via PepMap columns (150 µm × 15 cm, Thermo Fisher Scientific) in a trap and elution configuration. The mobile phases consisted of 0.1% formic acid in water (A) and 0.1% formic acid in 80% acetonitrile (B). The details of the LC gradient and flow rate for 100, 144, 180 and 300 SPD cycles are described in Supplemental Table 3.

**PRM Acquisition Method Development**

The list of 83 peptide sequences with specified charge states was imported into a Skyline and configured to have an iRT calculator with Pierce Peptide Retention Time Calibration™ (PRTC) standards, which included 14 more peptides in the assay whose relative retention times were known. The spectral library and iRT values for the 83 peptides were predicted via the Prosit inference engine built into Skyline, with the 30% normalized collision energy (NCE) parameter selected, and the top 15 transitions were retained. For each throughput, the PRM Conductor program was launched from Skyline and unscheduled targeted methods were created for each experimental sample-per-day (SPD) throughput (6).

The 83 heavy peptides were added to a vial at a concentration of 40 fmol/µl with PRTC at 50 fmol/µl. An unscheduled replicate injection was performed for each throughput, and the data were imported into separate Skyline documents. Skyline peak picking aided by the Prosit-predicted spectra and iRT values and was manually judged to be accurate for all the analytes via the retention time residual plot and the report table sorted by low-to-high spectral dot products to investigate any questionable precursors. This process was aided by the high degree of correspondence between the predicted and actual spectra and retention times. A further round of PRM Conductor was used to refine the methods. The “Keep all precs.” option was selected so that no precursors were removed, while transitions were filtered using the parameters Min. Abs. Area 1000, Min. Signal/Background 2.0, Min Rel. Area 0.05, Min. Time Corr. 0.8, and Min. Transitions 5. After accepting the picked LC peaks and using the Send to Skyline button to remove filtered transitions, the skyline documents were resaved, and the final spectral/iRT libraries were saved in skyline in the .blib format.

The acquisition parameters were Min. Dwell Time 5.0 milliseconds, Scan Rate 125 kDa/sec, and Acq. Window Opt., Maximum AGC Time Mode Dynamic, HCD activation with 35% NCE, Isolation Window 1.0 Th, and throughput-dependent parameters are listed in Supplemental Table 1. The Acq. Window Opt. selection slightly broadens the acquisition windows at the beginning and end of the assays. Notably, the optimized scan range used for the 300 SPD assay means that each peptide uses a customized analysis scan range that spans the m/z range of the accepted transitions. This mode offers an increased acquisition rate for potentially less ion injection time in the retention time regions of the assay where many precursors elute. However, precursors that elute in less crowded retention time regions may still be acquired with longer injection times because the Dynamic Maximum Injection Time mode distributes extra cycle time to the analysis of active precursors, as described previously (6). The Abs. Quan. The option was selected, which adds the light precursors to the exported instrument methods, making the final assays contain 83 light peptides, 83 heavy peptides, and 14 PRTC peptides for a total of 180 precursors.

The next experimental step was to confirm the retention times of the peptides in the presence of 300 ng of plasma matrix. Instrument methods were exported from the PRM Conductor, which contained an $MS^1$ experiment, a targeted $MS^2$ experiment describing the acquisition schedule of the analytes, and an additional adaptive RT DIA experiment to collect reference data. Because the neat peptide sample was significantly different from the plasma samples used for the bulk of the experiments, alignment from the neat sample to the plasma samples could not be directly performed. A replicate was acquired for each throughput using these conditions, with the results imported into separate Skyline documents. PRM Conductor was subsequently used on these replicates to create final methods using the parameters in Supplemental Table 1, but the instrument method template had the dynamic time scheduling option set to adaptive RT. In this case, the PRM Conductor creates instrument method files that embed the compressed reference spectra from the adaptive RT DIA experiment to be used for real-time updating of the acquisition schedule to account for the analyte retention time drift (7). The final spectral/RT libraries were saved in Skyline for the plasma matrix condition.

**Stellar Instrument Settings**

The retention times of the 83 heavy peptides and 14 PRTC peptides were determined using an instrument method with MS and $MS^2$ experiments. The MS experiment had a scan range of m/z 350--1250 and a scan rate of 125 kDa/sec. The RF Lens was 30%, AGC Target was Standard ($3e^4$) and Maximum Injection Time Mode was Auto. For the $MS^2$ experiment, the mass list was entered with the appropriate target m/z and z values, and Time Mode was set to Unscheduled. The Q1 isolation width was 1.0 Th, Activation Type was HCD with 30% normalized collision energy (NCE). The scan rate was 125 kDa/sec with Scan Range m/z 200--1500. The RF Lens was 30%, AGC Target was Standard ($1e^4$), and Maximum Injection Time Mode was Dynamic with 6 points per peak. The Chromatographic Peak Width was set to 10 seconds, and Collision Cell Gas Pressure was 8 mTorr. The data were imported into Skyline, where LC peaks were manually confirmed based on comparisons to *in silico* predictions of spectra and retention time using Prosit integrated into Skyline.

The retention times of the heavy labeled peptides in plasma were then determined with instrument methods that were largely the same except the number of points per peak was set to 7, and each throughput was given a customized Chromatographic Peak Width setting and Acquisition window width in PRM Conductor in the exported methods. The Chromatographic Peak Width in seconds was 6.85, 5.52, 4.78, and 4.5, and the nominal acquisition width in minutes was 2.0, 2.0, 1.5, and 0.5 for the respective throughputs of 100, 144, 180, and 300 samples per day. In PRM Conductor the Acquisition Window Optimization button was selected, which somewhat expands the Acquisition Windows especially at the start and end of the experiments. These instrument methods also included an Adaptive RT experiment that gathered reference data for the subsequent runs. PRM Conductor was used to create the instrument methods and Skyline analysis files, using Refine Targets settings of Min Abs. Area 400, Min Signal/Backgnd. 2.0, Min. Rel. Area 0.05, Min Time Corr. 0.8, Min Good Trans 3, and the Keep All Precs. option was selected so that no peptides were eliminated from the assay.

Final targeted methods were created for the 83 heavy labeled and 83 endogenous peptides plus the 14 PRTC peptides. Points per peak was 8, 8, 7, 7, the Chromatographic Peak Width in seconds was 7, 5, 5, and 4.5 and the nominal acquisition window width in minutes was 1.0, 0.6, 0.5, 0.2 for the respective throughputs of 100, 144, 180, and 300 samples per day. Only the 300 samples per day method utilized the Optimize Scan Range option in the PRM Conductor to increase the acquisition speed, while the other throughputs used the Scan Range of m/z 200-1500. These instrument methods included an Adaptive RT experiment with a reference file from the previous heavy-only plasma runs.

After analysis of the first dilution curves, 3 peptides were selected for the analysis via $MS^3$ for both heavy and light peptides instead of $MS^2$ due to the presence of interference in the endogenous peptide. $MS^2$ fragments were manually selected as $MS^3$ precursors for these peptides, although the PRM Conductor can also export methods with automatic $MS^3$ precursor selection. The $MS^3$ acquisitions used a Q1 isolation width of 2.0 instead of the 1.0 used for $MS^2$ analysis. For the following peptides, if resonance collision included dissociation (CID in the method editor) was used, the NCE was 30% for 10 msec for the first activation stage, and NCE was 35% for 5 msec on the second activation stage. The CID is applied serially to each $MS^3$ precursor, so the total acquisition time increases linearly with each precursor. The activation Q was 0.22. The collision cell collision-induced dissociation (HCD in the method editor) had NCE of 35%. The activation types used for each stage (first/second) for each protein:peptide was CLUS:EIQNAVNGVK (CID/HCD), CRP:ESDTSYVSLK (CID/CID), and CO9:TSNFNAAISLK (CID/HCD). The $MS^3$ Acquisition parameters are listed in Table 1.

**Table 1. $MS^3$ acquisition parameters**

| Peptide Sequence | Activation Type Stage 1 | Activation Energy Stage 1* | Activation Type Stage 2 | Activation Energy Stage 2* |
|---|---|---|---|---|
| EIQNAVNGK | CID | 30/10/0.22 | HCD | 35 |
| ESDTSYVSLK | CID | 30/10/0.22 | CID | 35/5/0.22 |
| TSNFNAAISLK | CID | 30/10/0.22 | HCD | 35 |

*Parameters for resonance CID activation are the normalized collision energy/activation time in milliseconds and the activation Mathieu q parameter. For HCD activation, the normalized collision energy is given.

**Loading Curve Analysis**

Loading curve experiments were conducted at 100, 144, 180, and 300 SPD using four independent, optimized, and scheduled PRM methods across 11 different SIL load levels: 120, 40, 13.3, 4.4, 1.48, 0.49, 0.16, 0.0548, 0.01829, 0.006097, and 0 fmol SIL on the column. A 300 ng digested control plasma pool was added as the matrix for all load levels. Five injections were performed for each loading level across all loading curves.

**Data analysis**

After the PRM data were acquired, the raw files from the PRM run were imported into Skyline Daily (version 24.1.1.254) for peak detection, integration, and visualization. The peaks were integrated with a tolerance of ±0.5 m/z. The area under the curve (AUC) was calculated for each peptide. The limits of detection and quantification were determined for each peptide and each PRM throughput method.

For the IBD samples, the peak area ratios (light to heavy standard) of the top three ions were exported from Skyline and used for the final statistical analysis. For all peptides, we employed the ratio dot product (rdopt) to assess the similarity between the heavy standard and light signals, flagging any samples with light signals that were too low for reliable detection. The threshold for reliable detection of a particular peptide in specific samples was set at a ratio dot product (rdotp) greater than 0.90.

**Skyline Curve Fitting**

The lower limits of quantification and detection (LLOQ and LLOD) were calculated with Skyline using traditional means, within Skyline ver. 24.1.1.202. The average LC peak area at each concentration level versus the spike-in concentration was modeled with a bilinear fit, where one line was constrained to slope = 0 and the second was constrained to intersect with the first. The LLOD was determined at the concentration where the two lines intersect. The LLOQ is determined by traversing the metrics for the coefficient of variation (CV) and model error at each level from high to low and selecting the LLOQ as the lowest concentration where the CV < 20% and the error < 20%.

**Statistical Analysis**

Considering the nature of repeated sample collection at different time points in the current study, the association between protein level and IBD diagnosis was performed using linear mixed model (LMM) after inverse rank normalization of the readout from the Stellar platform. Age and sex were included as covariates to control for potential confounding factor. Spearman correlation was performed to examine the correlation between clinically measured the CRP and CRP readout from Stellar. All the statistical analysis were performed using R 4.3.3 (www.r-project.org).

**CRP Antibody Based Clinical Assay**

Measurement of CRP in patient blood was performed by immunoturbidimetric assay on an Olympus system (Melville, N.Y.) as described previously (8).
